# A SoxB gene acts as an anterior gap gene and regulates posterior segment addition in the spider *Parasteatoda tepidariorum*

**DOI:** 10.1101/298448

**Authors:** Christian L. B. Paese, Anna Schoenauer, Daniel J. Leite, Steven Russell, Alistair P. McGregor

## Abstract

The Sox gene family encode a set of highly conserved HMG domain transcription factors that regulate many key processes during metazoan embryogenesis. In insects, the SoxB gene *Dichaete* is the only Sox gene known to be involved in embryonic segmentation. To determine if similar mechanisms are used in other arthropods, we investigated the role of Sox genes during segmentation in the spider *Parasteatoda tepidariorum*. While *Dichaete* does not appear to be involved in spider segmentation, RNAi knockdown of the closely related *Sox21b-1* gene results in a gap like phenotype in the developing prosoma and also perturbs the sequential addition of opisthosomal segments. We show that this is in part due to a role for *Sox21b-1* in regulating the expression of *Wnt8* and influencing Delta-Notch signalling during the formation of the segment addition zone. Thus, we have found that two different mechanisms for segmentation in a non-mandibulate arthropod are regulated by a Group B Sox gene. Our work provides new insights into the function of an important and conserved gene family across arthropods, and the evolution of the regulation of segmentation in these animals.

## Introduction

Arthropods are the most speciose and widespread of the animal phyla, and it is thought that their diversification and success is at least in part explained by their segmented body plan (1). In terms of development, insects utilise either derived long germ embryogenesis, where all body segments are made more or less simultaneously, or short/intermediate germ embryogenesis, where a few anterior segments are specified and posterior segments are added sequentially from a growth or segment addition zone (SAZ) (2, 3). It is thought that segmentation in the ancestral arthropod resembled the short germ mode seen in most insects (2, 4). Understanding the regulation of segmentation more widely across the arthropods is important for understanding both the development and evolution of these highly successful animals.

We have a detailed and growing understanding of the regulation of segmentation in various insects, especially the long germ dipteran *Drosophila melanogaster* and the short germ beetle *Tribolium castaneum*. However, studies of other arthropods including the myriapods *Strigamia maritima* and *Glomeris marginata*, and chelicerates, such as the spiders *Cupiennius salei* and *Parasteatoda tepidariorum*, have provided important mechanistic and evolutionary insights into arthropod segmentation (2, 5–9). Previous studies have shown that different genetic mechanisms are used to generate segments along the anterior-posterior axis of spider embryos. In the anterior tagma, the prosoma or cephalothorax, the cheliceral and pedipalpal segments are generated by dynamic waves of *hedgehog* (*hh*) and *orthodenticle* (*otd*) expression (10, 11). The leg bearing segments are specified by gap gene like functions of *hunchback* (*hb*) and *distal-less* (*dll*) (12, 13). In contrast, the segments of the posterior tagma, the opisthosoma or abdomen, are generated sequentially from a SAZ. This process is regulated by dynamic interactions between Delta-Notch and Wnt8 signalling to regulate *caudal* (*cad*), which in turn is required for oscillatory expression of pair-rule gene orthologues including *even-skipped* (*eve*), and *runt* (*run*) (4, 14, 15). Interestingly, these pair-rule gene orthologues are not involved in the production of the prosomal segments (15). Therefore, the genetic regulation of segmentation along the anterior-posterior axis in the spider exhibits similarities and differences to segmentation in both long germ and short germ insects.

The Group B Sox family gene *Dichaete* is required for correct embryonic segmentation in the long germ *D. melanogaster*, where it regulates pair-rule gene expression (16,17). Interestingly, it was recently discovered that a *Dichaete* orthologue is also involved in segmentation in the short germ *T. castaneum* (18). This similarity is consistent with work inferring that these modes of segmentation are more similar than previously thought and provides insights into how the long germ mode evolved (18, 19, 20). However, it appears that despite these similarities, *Dichaete* can play different roles in *D. melanogaster* and *T. castaneum* consistent with the generation of segments simultaneously via a gap gene mechanism in the former and sequentially from a posterior SAZ in the latter (18).

We recently described the discovery of 14 Sox genes in the genome of the spider *P. tepidariorum* (21) and that several of the spider Sox genes are represented by multiple copies likely produced during the whole genome duplication (WGD) in the lineage leading to this arachnid (21). Interestingly, while *Dichaete* is not expressed in a pattern consistent with a role in segmentation (21), we found that the closely related SoxB gene, *Sox21b-1*, is expressed in both the prosoma and opisthosoma before and during segmentation (21). Here we report that in *P. tepidariorum*, *Sox21b-1*, regulates both prosomal and opisthosomal segmentation. In the prosoma, *Sox21b-1* has a gap-like gene function and is required for the generation of the four leg bearing segments. In addition, *Sox21b-1* appears to act upstream of both *Delta* and *Wnt8* to regulate the formation of the SAZ, and knockdown of *Sox21b-1* results in truncated embryos missing all opisthosomal segments. Therefore, while prosomal and opisthosomal segments are generated by different mechanisms in the spider, our analysis shows that *Sox21b-1* is required for segmentation in both regions of the developing spider embryo.

## RESULTS

### *Sox21b-1* is maternally deposited and is subsequently expressed in the germ disc and germ band of spider embryos

We previously identified and assayed the expression of the complement of Sox genes in the genome of the spider *P. tepidariorum* (21). Our phylogenetic analysis indicates that *P. tepidariorum Sox21b-1* and its paralog *Sox21b-2* are members of the Sox group B, closely related to the *Drosophila Dichaete* and *Sox21b* genes (Fig. S1). In insects (22, 23), *Dichaete, Sox21a* and *Sox21b* are clustered in the genome, however, both *Sox21b* paralogs are dispersed in the spider genome (21). This suggests that *Sox21b-1* and *Sox21b-2* possibly arose from the WGD event in the ancestor of arachnopulmonates (24), rather than by a more recent tandem duplication (20) (Fig. S1).

In light of its interesting expression pattern we elected to analyse *Sox21b-1* further. Pre-vitellogenic *P. tepidariorum* oocytes contain a Balbiani’s body (25), where maternally deposited factors are enclosed, and we found that *Sox21b-1* is abundant in this region, indicating that it is maternally contributed (Fig. 1A). However, after fertilization *Sox21b-1* is not expressed again until early stage 5, when weak expression is detected throughout the germ disc, with stronger expression in more central cells (Fig. 1B). At late stage 5, expression becomes more restricted to the centre of the germ disc (Fig. 1C). During stages 5 and 6, the cumulus migrates to the rim of the germ disc, opening the dorsal field and giving rise to an axially symmetric germ band (Fig. 1D) (see 26). In early stage 6 embryos, *Sox21b-1* is observed in the middle of the presumptive prosoma in a broad stripe (Fig. 1D), which develops further during stage 7 in the region where the leg bearing segments will form (Fig. 1E). This expression pattern resembles the previously described expression of the gap gene *hb* (13).

**Figure 1.**
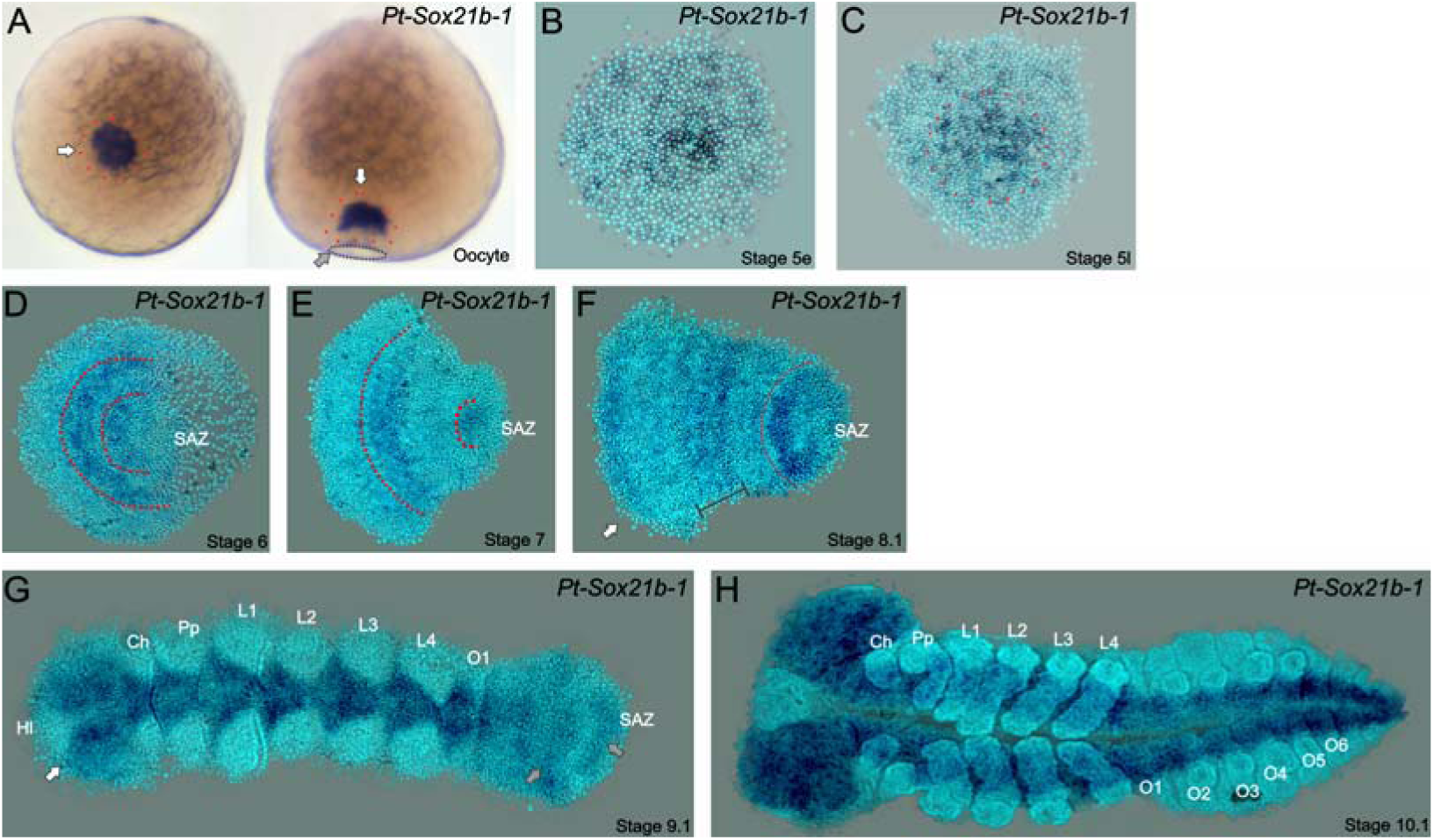
Expression of *Sox21b-1* in *P. tepidariorum* oocytes and embryos. **A)** Dorsal (left) and lateral (right) views of pre-vitellogenic oocytes showing *Sox21b-1* mRNA in the Balbiani’s body (red dashed circle and white arrows). The sperm implantation groove is indicated by a black dashed circle and grey arrow. **B**) At early stage 5, the expression of *Sox21b-1* appears in a salt and pepper pattern in the germ disc. **C**) Expression in the cumulus becomes stronger at late stage 5, with less expression at the periphery of the germ disc (dashed red circle). **D**) At stage 6 *Sox21b-1* is expressed in a broad stripe in the anterior (between the red dashed lines). **E**) At stage 7 there is expression in the region of the presumptive leg bearing segments and in the SAZ (both indicated by red dashed lines). **F**) At stage 8.1 *Sox21b-1* is still expressed in the SAZ and the presumptive leg bearing segments, but nascent expression is observed at the anterior of the germ band (indicated by the white arrows and black brackets). **G**) At stage 9.1, when the limb buds are visible the expression of *Sox21b-1* becomes restricted to the ventral nerve cord (anterior white arrow) and can be observed in the SAZ (posterior grey arrows). **H**) At stage 10.1, *Sox21b-1* expression is restricted to the ventral nerve cord and the head lobes. Ch: Chelicerae; HL: Head lobes; L1 to L4: Prosomal leg bearing segments 1 to 4; O1 to O6: Opisthosomal segments 1 to 6; SAZ: Segment addition zone. Ventral views are shown with anterior to the left, except as described for oocytes.

During these and subsequent stages, dynamic expression of *Sox21b-1* is observed in the SAZ and the most anterior region of the germ band that will give rise to the head segments (Fig. 1F-H). Later in development, the expression of *Sox21b-1* resembles that of *SoxNeuro* (*SoxN*), another Group B Sox gene (21). This expression is similar to that of both *SoxN* and *Dichaete* in *D. melanogaster*, which are expressed in neuroblasts of the neuroectoderm and then differentiating neurons in the ventral nerve cord (22) (Fig. 1G, H). Expression of the related group B Sox genes, *Dichaete* and *Sox21b-2* are not detected in *P. tepidariorum* during embryonic development (20). The expression of *Sox21b-1* in the embryo suggests that it is involved in both anterior and posterior segmentation in this spider, as well as later during nervous system development.

### *Sox21b-1* regulates prosomal and opisthosomal segmentation

To assay the function of *Sox21b-1* during embryogenesis we knocked down the expression of the gene using a parental RNAi approach (27). We observed three phenotypic classes, which were consistent between both of the non-overlapping *Sox21b-1* fragments we used for RNAi (Fig. 2; Fig. S2; Table S2).

**Figure 2.**
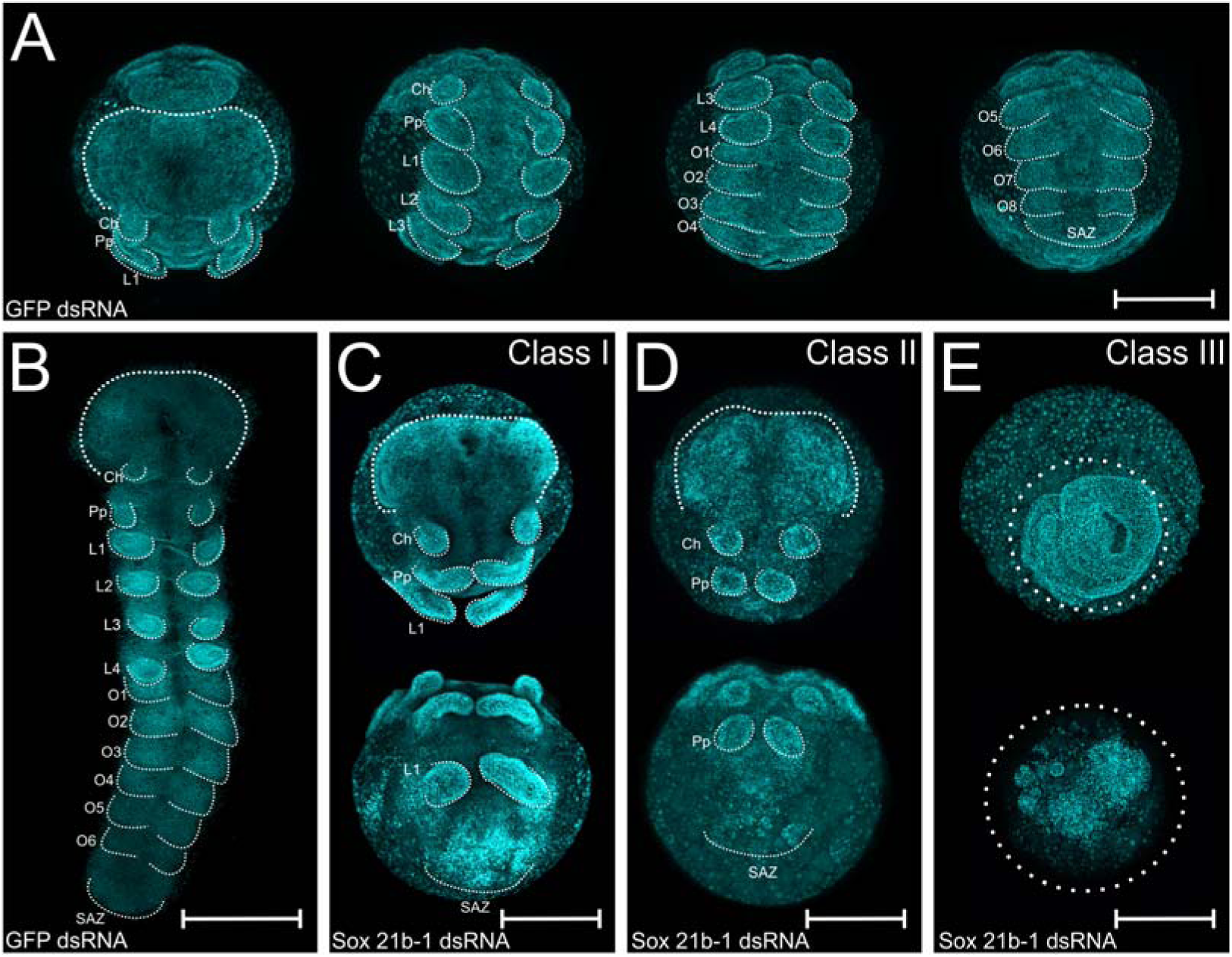
Embryo phenotypes after *Sox21b-1* parental RNAi knockdown. Whole mount (**A**) and flat mount (**B**) control embryos at stage 9 stained with DAPI. Stage 9, Class I (**C**), Class II (**D**) and Class III (**E**) phenotypes from *Sox21b-1* knockdown. In the control embryos (A and B), the head, cheliceral (Ch), pedipalpal (Pp), prosomal walking limbs (L1 to L4), opisthosomal segments (O1 to O6) and a posterior SAZ are all clearly visible as indicated. **C**) Class I phenotype embryos show a morphologically normal head, pairs of chelicerae, pedipalps and first walking limbs (Ch, Pp, L1), but a disorganised cluster of cells in the posterior where L2-L4, opisthosomal segments and the SAZ should be. **D**) Class II phenotype embryos consist of fewer cells, but still form a head, chelicerae, pedipalps (Ch, Pp) and a structure resembling the SAZ in the posterior. **E**) Class III embryos exhibit the most severe phenotype, where, after the germ disc stage, the embryo fails to form an organised germ band. Anterior is to the top, scale bars: 150 µm.

Class I embryos developed a presumptive head region (Fig. 2A-C), as well as normal cheliceral, pedipalpal and first leg bearing (L1) segments (Fig. 2C). The identity of these segments was confirmed by expression of *labial* (*lab*) in the pedipalps and L1, and *Deformed*-*A* (*Dfd-*A) in L1 (Fig S3A-B). However, the other three leg bearing segments, L2 - L4, as well as all of the opisthosomal segments were missing in Class I embryos. These embryos exhibited a truncated germ band, terminating in disorganised tissue in the region of the SAZ (Fig. 2C). In the case of Class II phenotypes, embryos only differentiated the head region and the cheliceral and pedipalpal segments (Fig. 2D; Fig S3A-B): all leg bearing segments of the prosoma and opisthosomal segments produced from the SAZ were missing (Fig. 2D). In Class III embryos, the germ band did not form properly from the germ disc (Fig. 2E) and we therefore looked earlier in development to understand how this severe phenotype arose. We observed that the formation of the primary thickening occurs normally at stage 4 (27, 28, 29), but subsequently the cumulus, the group of mesenchymal cells that arise as the primary thickening at the centre of the germ disc, fails to migrate properly to the rim of the germ disc during stage 5 (Fig. S4). Since migration of the cumulus is required for the transition from germ disc to germ band, this observation at least in part explains the subsequent Class III phenotype. In some embryos, we did observe the opening of the dorsal field in stage 6 embryos: therefore, we suggest these embryos later develop Class I and II phenotypes (Fig. S4B-C).

We next examined the effect of *Sox21b-1* depletion on cell death and proliferation at stages 5 and 9 in knockdown and control embryos using antibodies against Caspase-3 and phosphorylated Histone 3 (PHH3) (Fig. S5). At the germ disc stage there is no detectable cell death in control embryos (n = 10), but we observed some small clusters of apoptotic cells in the *Sox21b-1* knockdown embryos (n = 10) (Fig. S5A-B). At stage 9, a few cells expressed Caspase-3 in the posterior-most part of the SAZ (Fig. S5C), but we did not observe cell death in this region of *Sox21b-1* knockdown embryos (Fig. S5D). However, we did observe pronounced cell death in the anterior extraembryonic layer of the same embryos, (n = 10) (Fig. S5D).

Expression of PHH3 at stages 5 and 9, indicated that *Sox21b-1* knockdown embryos show decreased cell proliferation compared to controls (n= 10 for each) (Fig. S5E-H). Interestingly the cells were also clearly larger in *Sox21b-1* knockdown embryos compared to controls, which may reflect perturbed cell proliferation (Fig. S5E-H). Thus, our functional analysis shows that *Sox21b-1* regulates cell proliferation and the transition from radial to axial symmetry. Moreover, *Sox21b-1* is involved in two different segmentation mechanisms in spiders: it has a gap gene like function in the prosoma, as well as a requirement for the formation of the SAZ and subsequent production of opisthosomal segments.

### Effects of *Sox21b-1* knockdown on the germ disc and mesoderm

In *P. tepidariorum, decapentaplegic* (*dpp*) and *Ets4* are required for cumulus formation (29, 30). To investigate if *Sox21b-1* is involved in the formation of this cell cluster we assayed the expression of *dpp* and *ets4* in *Sox21b-1* RNAi knockdown embryos. However, both genes were expressed normally and cumulus formation was unaffected (Fig. 3 E, F).

**Figure 3.**
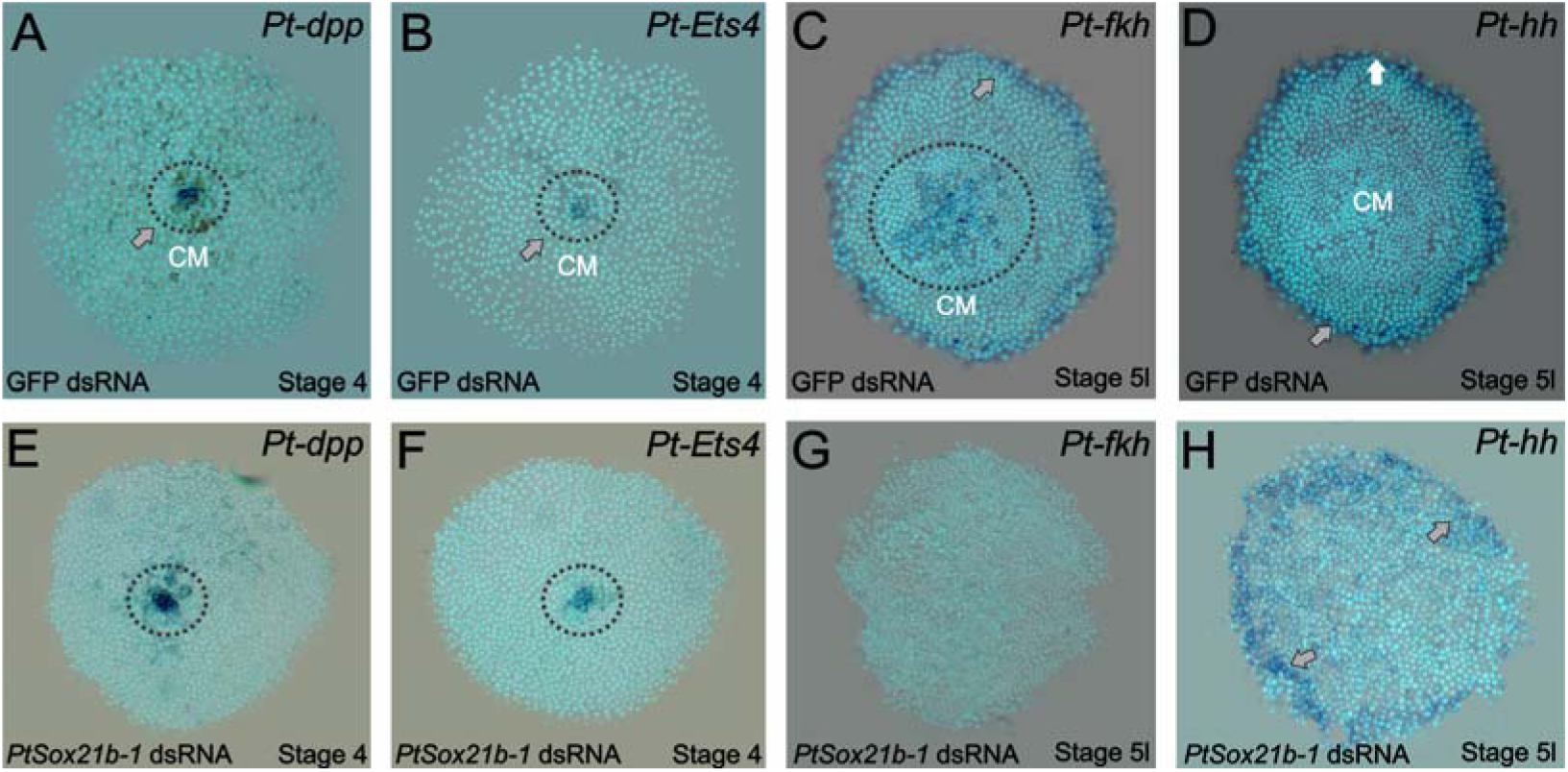
Gene expression in control and *Sox21b-1* knockdowns at the germ disc stage. *Pt-dpp* (A) and *Pt-Ets4* (B) are expressed in the forming cumulus (CM) in the centre of the germ disc at stage 4 (grey arrow and dotted circle). This expression is unaffected by knockdown of *Sox21b-1* (**E** and **F**) (n= 30 for each gene). **C**) *Pt-fkh* is expressed at the rim and centre of the germ disc at late stage 5 (grey arrow and dotted circle in C), but expression is lost in *Sox21b-1* embryos (n= 30) (**G**). *Pt-hh* expression at the rim of the germ disc (**D**) is normal in *Sox21b-1* knockdown embryos (**H**) (grey arrows).

The rim of the spider germ disc develops into the head structures and is regulated in part by *hh*, while the mesodermal and endodermal layers of the head are specified by the mesendodermal gene *forkhead* (*fkh*) (10, 27). To investigate if anterior expression of *Sox21b-1* (Fig. 1B) is involved in the formation of the head rudiment and differentiation of the mesodermal and endodermal layers in particular, we assayed the expression of *hh* and *fkh* in class I and II *Sox21b-1* knockdown embryos.

*hh* is expressed at the rim of the germ disc in the ectoderm (Fig. 3 D) (31) and remains unaffected by *Sox21b-1* knockdown (Fig. 3H). *fkh* is also expressed in cells around the rim, as well as in the centre of the germ disc in mesendodermal cells (Fig. 3C). In *Sox21b-1* knockdown embryos both of these *fkh* expression domains are lost (Fig. 3G) and it therefore appears that *Sox21b-1* is required for specification of mesendodermal cells in the germ disc of spider embryos. Indeed in the germ disc at stage 5, when *fkh* expression commences, we observed invaginating cells forming a second layer (Fig. S3G), however, in *Sox21b-1* knockdown embryos we observed a lower number of invaginating cells, which exhibit bigger nuclei compared to the controls (Fig. S3H).

In both spiders and flies, the *twist* (*twi*) gene is involved in mesoderm specification (32) and we therefore examined the expression of this gene after *Sox21b-1* knockdown to further evaluate if the loss of *fkh* affects the formation of the internal layers. *twi* is expressed in the visceral mesoderm of the limb buds from L1 to L4, in the opisthosomal segments O1 to O4 and in an anterior mesodermal patch in the central part of the developing head (32) (Fig. S3E). While the head expression persists in *Sox21b-1* class I embryos, expression in all the limb and opisthosomal segments is lower or absent (Fig. S3F). In orthogonal projections the anterior-most region of the embryo three layers of cells can be identified in control embryos (Fig. S3I). However, in *Sox21b-1* knockdown embryos the formation of these layers is perturbed (Fig. S3J). These data suggest that the ectodermal segmentation in the prosomal region occurs even upon a reduction of the internal layers of the embryo.

### Effects of *Sox21b-1* knockdown on segmentation

In *P. tepidariorum*, formation of the SAZ and production of posterior segments requires the Wnt8 and Delta-Notch signalling pathways (14, 15). Interactions between these pathways regulate *hairy* (*h*) and, via *cad*, the expression of pair-rule gene orthologues including *eve* (14, 15). To better understand the loss of segments we observe in *Sox21b-1* knockdown embryos we analysed the expression of *Dl, Wnt8, h* and *cad* in these embryos compared to controls.

*Dl* is expressed at stages 5 and 6 in the forming SAZ, in the region of the L4 primordia, and in the presumptive head (33) (Fig. 4A). Subsequently at stage 9, *Dl* expression is visible in clusters of differentiating neuronal cells and oscillates in the SAZ, an expression pattern associated with the sequential addition of new segments (Fig. 4B). In *Sox21b-1* knockdown embryos, *Dl* expression is not detected at stage 5 (Fig. 4C) and is absent in the posterior at stage 9 (Fig. 4D). However, expression in the anterior neuroectoderm seems normal up to the pedipalpal segment, although neurogenesis is apparently perturbed in the presumptive L1 segment (Fig. 4D). This suggests that the ectoderm up to the L1 segment differentiates normally, but the development of the SAZ and posterior segment addition controlled by *Dl* is lost upon *Sox21b-1* knockdown.

**Figure 4.**
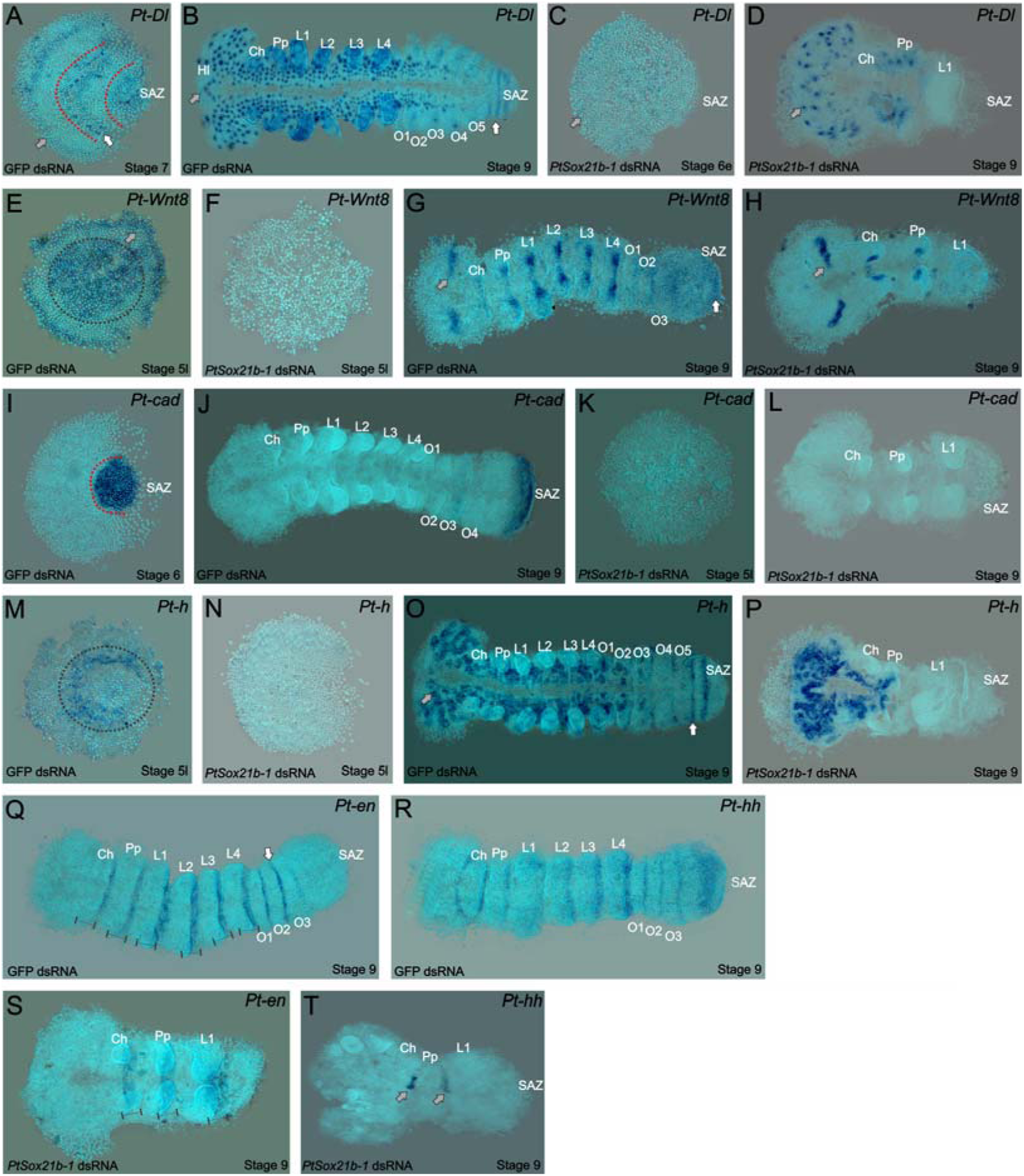
Expression of segmentation genes in *Sox21b-1* pRNAi embryos. **A** and **B**) *Pt-Dl* expression at late stage 6/early stage 7 is dynamic in the SAZ and is also observed in the presumptive head region and prosoma of the embryo (red dotted lines and grey arrows). **B**) At stage 9, *Pt-Dl* expression is seen in the SAZ (white arrow) but is restricted to the clusters of proneural differentiation in the anterior region of the embryo (grey arrow in the head lobes). **C**) In *Sox21b-1* knockdown embryos, *Pt-Dl* expression is not detectable in late stage 5/early stage 6 embryos (grey arrow) but can still be observed in the anterior ventral neuroectoderm at stage 9 up to the pedipalpal segment (n= 17 and n = 14 for stage 5 and 9, respectively) (**D**). *Pt-h* expression at stage 5 in control embryos is seen at the rim and in the centre of the germ disc (black dotted circle in **E**), which is lost in *Sox21b-1* knockdown embryos (**F**). At stage 9, *Pt-h* expression resembles *Pt-Dl*, both in the control and *Sox21b-1* knockdown embryos (**G** and **H**) (n= 15 for both stages). *Pt-Wnt8* expression is similar to *Pt-h* in stage 5 control embryos (black dotted circle in the centre, grey arrow to the rim) and is also lost in *Sox21b-1* knockdown embryos (n= 11) (**I** and **J**). Control embryos at stage 9 show the expression of *Pt-Wnt8* in the medial region of the head (grey arrow), and in distal parts of each segment up to the SAZ (white arrow) (**K**). In *Sox21b-1* knockdown embryos at the same stage, the brain (grey arrow), cheliceral and pedipalpal expression is still present, but the posterior expression is lost (n= 17 for each stage) (**L**). *Pt-cad* is expressed in the SAZ at late stage 5/early stage 6 embryos (**M**), which persists throughout to stage 9 control embryos (**N**). However, *Pt-cad* expression is lost upon *Sox21b-1* knockdown (n= 20 for each stage) (**O** and **P**). *Pt-en* expression is present in the posterior of each segment (black lines in **Q**), and in cheliceral, pedipalpal and L1 segments in *Sox21b-1* knockdown embryos at stage 9 (n= 10) (**S**). *Pt-hh* expression in control embryos at stage 9 is seen in the posterior of each segment and in the SAZ (**R**). When *Sox21b-1* is knocked-down, *Pt-hh* embryos show expression in the middle posterior of the cheliceral and pedipalpal segments (n= 8) (**T**). Ch: Chelicerae; HL: Head Lobes; L1 to L4: Prosomal leg bearing segments; O1 to O5: Opisthosomal segments; SAZ: Segment Addition Zone. Anterior is to the left in stage 9 embryos.

*Wnt8* is initially expressed at stage 5 in the centre and at the rim of the germ disc (Fig. 4E). At stage 9, striped expression of *Wnt8* is seen from the head to the posterior segments and in the posterior cells of the SAZ (Fig. 4G). Knockdown of *Sox21b-1* results in the loss of *Wnt8* expression in late stage 5 embryos (Fig. 4F). At stage 9, *Wnt8* expression is observed in the cheliceral, pedipalpal and first walking limb segments of *Sox21b-1* knockdown embryos, but no expression is detected in the remaining posterior cells (Fig. 4H). Consistent with the loss of *Dl* and *Wnt8*, *cad* expression is also lost in stage 5 and stage 9 *Sox21b-1* knockdown embryos (Fig. 4I-L). These observations indicate that *Sox21b-1* is acts upstream of Wnt8 and Delta-Notch signalling to regulate the formation of the SAZ and the subsequent production of posterior segments. In support of this regulatory relationship we find that *Sox21b-1* expression is still detected in the posterior regions of the truncated embryos produced by RNAi knockdown of either *Dl* or *Wnt8* (Fig. 5).

**Figure 5.**
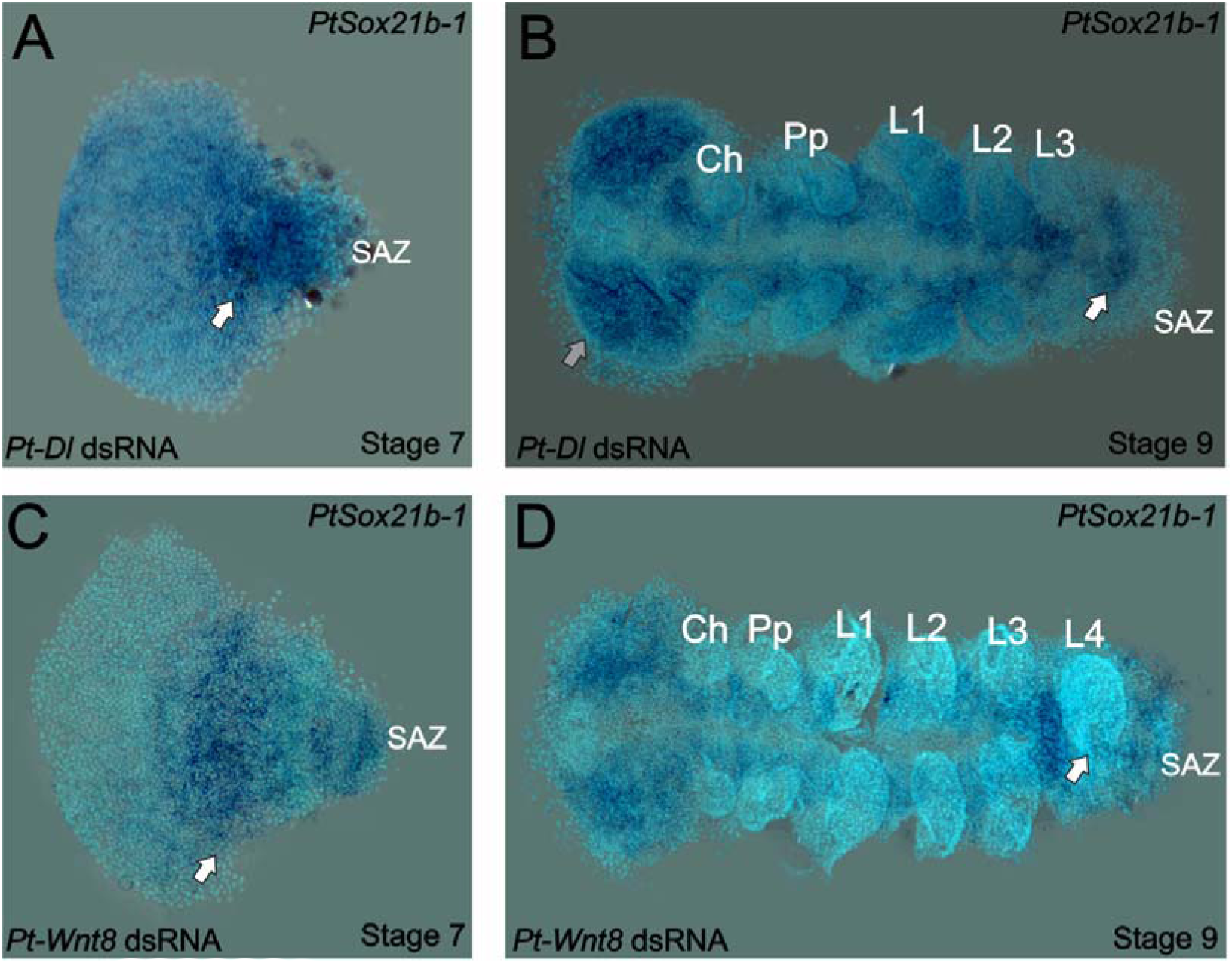
Expression of *Sox21b-1* in *Dl* and *Wnt8* pRNAi embryos. Ventral view of stage 7 and 9 knockdown embryos for *Pt-Dl* (**A** and **B**) and *Pt-Wnt8* (**C** and **D**). In knockdown embryos for both *Pt-Dl* and *Pt-Wnt8*, *Sox21b-1* is still expressed at the germ band stage (**A** and **C**), in a dynamic pattern in the remaining SAZ cells, and in the forming segments in the presumptive prosoma of the embryo (white arrows). In stage 9 *Pt-Wnt8* knockdown embryos, *Sox21b-1* remains highly expressed in the ventral nerve cord (**D**). *Pt-Dl* knockdown embryos lack the posterior L4 segment (white arrow), but brain formation appears normal (grey arrow) (**B**). *Pt-Wnt8* embryos show a fusion of the L4 limb buds, and *Sox21b-1* is still expressed in the remaining SAZ cells (**D**). Anterior is to the left in all panels.

The spider orthologue of *h* is expressed in the presumptive L2-L4 segments and dynamically in the SAZ (14) (Fig. 4M, O). In late stage 5 *Sox21b-1* knockdown embryos, the expression of *h* is lost throughout the entire germ disc (Fig. 4N). In addition, in Class I phenotype embryos at stage 9, the expression of *h* is completely absent in the tissue posterior to the pedipalpal segment (Fig. 4P). Therefore the loss of *h* expression is consistent with the loss of leg bearing segments in the anterior gap-like phenotype that results from knockdown of *Sox21b-1* as well as loss of segments produced by the SAZ.

To look at the effect of *Sox21b-1* knockdown on segmentation in more detail we examined the expression of *engrailed* (*en*) and *hh*. At stage 9 *en* is expressed segmentally from the cheliceral to the O3 segment in control embryos (Fig. 4Q). However, in *Sox21b-1* knockdown embryos, expression of *en* was only observed in the cheliceral, pedipalpal and L1 segments, consistent with the loss of all the more posterior segments (Fig. 4S). *hh* has a similar expression pattern to *en* at stage 9, except it exhibits an anterior splitting wave in the cheliceral segment and is also expressed earlier in opisthosomal segments and in the SAZ (Fig. 4R). Upon *Sox21b-1* knockdown, *hh* is only detected in shortened stripes in the cheliceral and pedipalpal segments (Fig. 4T).

Taken together, our analysis of *P. tepidariorum* embryos where *Sox21b-1* is depleted by parental RNAi reveals an important role for this Group B Sox gene in both gap-like segmentation of the prosoma, as well as posterior segment formation from the SAZ. These experiments further emphasise the critical role this class of transcription factors play in arthropod segmentation.

## Discussion

### A SoxB gene is required for two different mechanisms of spider segmentation

The Sox (Sry-Related High-Mobility Group box) gene family encodes transcription factors that regulate many important processes underlying the embryonic development of metazoans (34-37). One such gene, *Dichaete*, is expressed in a gap gene pattern and is involved in regulating the canonical segmentation cascade in *D. melanogaster* (16, 17). Recently, the analysis of the expression of *Dichaete* in the flour beetle *T. castaneum* strongly suggests a role in short germ segmentation (18), further supported by knockdown of the *Dichaete* orthologue in *Bombyx mori*, which resulted in the loss of posterior segmentation (38).

Here we show that, while *Dichaete* is not involved in spider segmentation (21), the closely related SoxB gene, *Sox21b-1*, regulates formation of both prosomal and opisthosomal segments. In the prosoma *Sox21b-1* has a gap gene role and is required for the specification of L1-L4 segments (Fig. 6), resembling the roles of *hb* and *Dll* in prosomal segmentation in this spider (12, 13) and, at least superficially, gap gene function in *Drosophila*.

**Figure 6.**
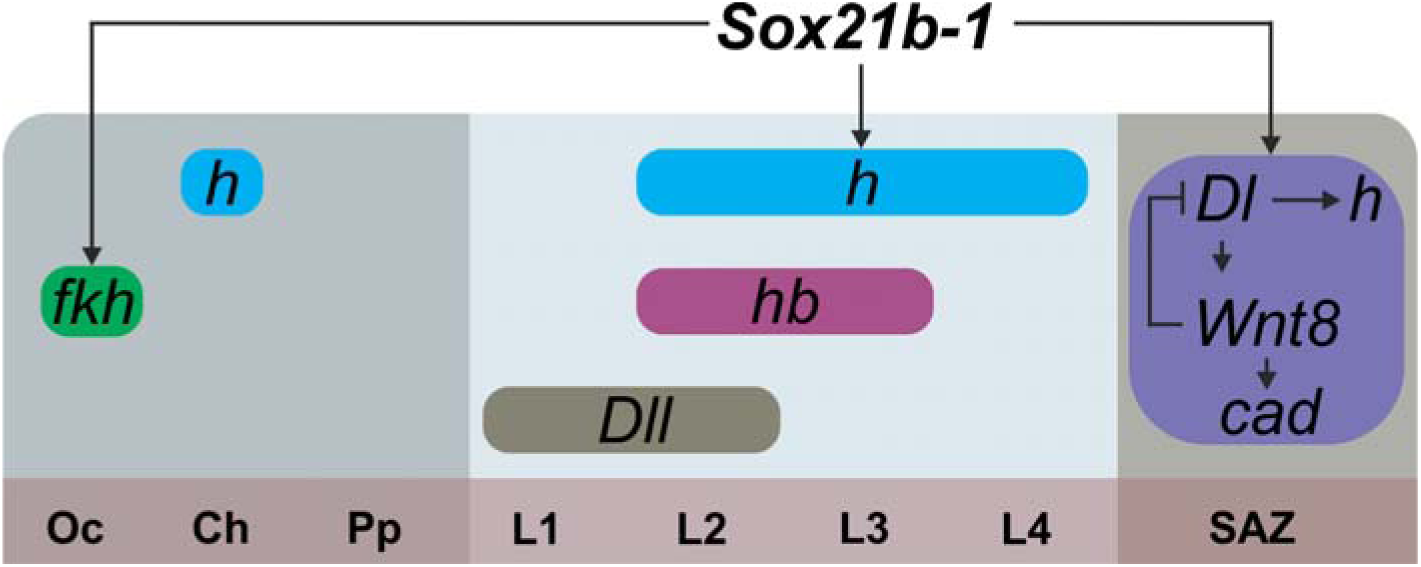
Summary of the regulation of spider segmentation. The interaction of *Sox21b-1* is presented in relation to genes involved in spider embryogenesis. We found that *fkh* expression requires *Sox21b-1* in the most anterior part of the head (OC, Ch, Pp segments). In the prosoma *Sox21b-1* has a gap gene like function and is required for the expression of *hairy*, while *Distal-less* (12) and *hunchback* (11) also act as gap genes during prosomal segmentation (L1-L4). The molecular control of segmentation in the SAZ involves a feedback loop between *Dl* and *Wnt8*, which acts upstream of *cad* and also controls the dynamic expression of *hairy* (15). We can infer from our results that *Sox21b-1* acts upstream of these genes in the SAZ.

In *Drosophila* the gap genes regulate pair rule gene expression, and while our results indicate that *Sox21b-1* is required for the expression of *h* and the generation of leg bearing prosomal segments (Fig 4E; Fig. 6), in contrast to insects, in spiders this does not involve the orthologues of *eve* and *runt* because they are not expressed in the developing prosomal segments (15, 39).

In the posterior, *Sox21b-1* knockdown perturbs SAZ formation and consequently results in truncated embryos missing all opisthosomal segments. Therefore, *Sox21b-1* regulates development of the SAZ, and our observations indicate this is at least in part through roles in organising the germ layers and specification of mesendodermal cells during stages 5 and 6. This is supported by the loss of *fkh* expression upon *Sox21b-1* knockdown, which is required for mesoderm and endoderm formation in both spiders and insects (10, 40, 41). Moreover, the subsequent dynamic expression of *Sox21b-1* in the SAZ after stage 6 is suggestive of a role in segment addition.

Our work on *Sox21b-1* provides an important new insight into the gene regulatory network (GRN) underlying the formation of the SAZ and the sequential addition of segments from this tissue. We show that *Sox21b-1* acts upstream of Wnt8 and Delta-Notch signalling in this GRN and is required for the activation of these important signalling pathways during posterior development (Fig. 6). Further work is needed to determine if Group B Sox genes, such as *Dichaete* and *Sox21b-1*, occupy a similar position in the regulatory hierarchy for posterior segmentation in other arthropods. This could provide important new insights into the evolution of the regulation of segmentation in arthropods since a Wnt-Delta-Notch-Cad regulatory cassette was probably used ancestrally in arthropods to regulate posterior development (4, 8, 9, 14, 15, 42). Interestingly, SoxB genes also cooperate with Wnt and Delta-Notch signalling in various aspects of vertebrate development including the patterning of neural progenitors and maintenance of the stem state in the neuroepithelium (35, 43, 44, 45).

### Sox21b-1 exhibits highly pleiotropic phenotypes during early spider embryogenesis

Our study shows that *Sox21b-1* is not only involved in segmentation but is maternally supplied and regulates cell division in the early germ disc, as well as the transition from radial to axial symmetry during germ band formation. Further experiments with *Sox21b-1* are required to fully elucidate the mechanisms by which it affects these early functions. Furthermore, while spider head development is less affect than trunk segmentation by knockdown of *Sox21b-1*, it is clear from our experiment that *Sox21b-1* regulates cell fate in this region. Interestingly, Sox2 is involved with the neuro-mesodermal fate choice in mice and *Dichaete* has a role in embryonic brain development in *Drosophila* (45, 46): consequently, SoxB genes may play an ancestral role in the patterning of the head ectoderm and mesoderm in metazoans (45, 46).

### The evolution of Sox21b-1

The evolution and diversification of Group B Sox genes in insects is not fully resolved due to difficulties in clearly assigning orthologues based on the highly conserved HMG domain sequence (17, 35, 47). However, despite these ambiguities it is clear that the *Dichaete* and *Sox21b* class genes in all arthropods examined to date are closely related and likely arose from a duplication in the common ancestor of this phylum (see 47 for discussion). Note that in insects *Dichaete*, *Sox21a* and *Sox21b* are clustered (29), however, while *Dichaete* and *Sox21a* are also clustered in *P. tepidariorum*, the *Sox21b* paralogs are dispersed in the genome of this spider (21). We believe it is highly significant that two very closely related SoxB genes are involved in segmentation in both the spider *P. tepidariorum* and in insects, pointing to an ancient role for this subfamily of Sox genes in invertebrates. Given the close similarity between the HMG domains of Sox21b and Dichaete, it is possible that in some lineages the Dichaete orthologue assumed the segmentation role, whereas in others it was Sox21b. In spiders, *Wnt8* is involved in posterior development while in other arthropods this role is played by *Wnt1/wg* (14), and therefore the evolution of *Sox21b-1* may have led to the co-option to different genes and developmental systems drift of the GRN for posterior development.

The spider contains an additional related SoxB gene, *Sox21b-*2, that possibly arose as part of the whole genome duplication event in the ancestor of arachnopulmonates over 400 million years ago (22). It will be interesting to examine any segmentation roles in other spiders and arachnids, including those that did not undergo a genome duplication. Finally, Blast searches of the Tardigrade *Hypsibius dujardini* genome reveal a single *Dichaete/Sox21b* class gene and it will be of some interest to characterise the expression and/or function of this gene in this sister group to the arthropods.

## Materials and Methods

### Spider Culture

*P. tepidariorum* were cultured at 25°C at Oxford Brookes University. The spiders were fed with *D. melanogaster* with vestigial wings and subsequently small crickets (*Gryllus bimaculatus*). Cocoons from mated females were removed and a small number of embryos were immersed in halocarbon oil for staging according to (26).

### Phylogenetic analysis of *P. tepidariorum* Sox genes

To identify the phylogenetic relationship of *P. tepidariorum* Sox genes the HMG domains of *Anopheles gambiae*, *Mus musculus*, *D. melanogaster*, *P. tepidariorum* and *S. mimosarum* Sox genes were aligned with ClustalW (21, 48). Phylogenetic analysis was performed in RAxML, with support levels estimated implementing the rapid bootstrap algorithm (1000 replicates) (49), under the PROTGAMMALG model of amino acid substitution, which was identified as best fitting using a custom Perl script from the Exelixis Lab website (https://sco.h-its.org/exelixis/web/software/raxml/hands_on.html).

### Fixation and gene expression analysis

Embryos ranging from the 1-cell stage to stage 13 were dechorionated and fixed according to (31) with a longer fixation time of 1 hr to facilitate yolk removal for flat-mounting. For immunohistochemistry, methanol steps were omitted. Ovaries from adult females were dissected in 1x PBS and fixed in 4% formaldehyde for 30 min. Probe synthesis and RNA *in situ* hybridisation was carried out with minor modifications to (27), omitting Proteinase K treatment and post-fixation steps. Poly-L-lysine (Sigma-Aldrich) coated coverslips were used for flat-mounting embryos. Nuclei were stained by incubating embryos in 1 μg/ml 4-6-diamidino-2-phenylindol (DAPI) in PBS with 0.1% Tween-20 for 15 min.

### Imaging, Live Imaging and Image Analysis

For imaging of flat-mounted embryos after *in situ* hybridisation, an AxioZoom V16 stereomicroscope (Zeiss) equipped with an Axiocam 506-Mono and a colour digital camera were used. Immunostained embryos were imaged with Zeiss LSM 800 or 880 with Airyscan confocal microscopes. For live imaging, embryos were aligned on heptane glue coated coverslips and submersed in a thin layer of halocarbon oil. Bright-field live imaging was performed using an AxioZoom V16 stereomicroscope, while fluorescence live imaging was performed with confocal microscopes. Image stacks were processed in Fiji (50) and Helicon Focus (HeliconSoft). Image brightness and intensity was adjusted in Corel PhotoPaint X5 (CorelDraw) and Fiji.

### Gene Isolation from cDNA

Fragment of genes were amplified using PCR and cloned into pCR4-TOPO (Invitrogen, Life Technologies). Oligonucleotide sequences are listed in Table S1.

### Immunohistochemistry

Immunostaining was carried out following (51) with minor modifications: antibodies were not pre-absorbed prior to incubation and the concentration of Triton was increased to 0.1%. The following primary antibodies were used: mouse anti-α-Tubulin DM1a (Sigma) (1:50), rabbit α cleaved caspase 3 (Cell Signaling - 9661) (1:200) and rabbit Anti-phospho-Histone H3 (Ser10) (Merck Millipore - 06-570). For detection the following secondary antibodies were used: donkey anti-mouse IgG Alexa Fluor 555 (Invitrogen) and goat anti-rabbit Alexa Fluor 647 (Invitrogen). The counterstaining was carried out by incubation in 1 μg/ml 4-6-diamidino-2-phenylindol (DAPI) in PBS + Triton 0,1% for 20 minutes.

### dsRNA synthesis and Parental RNA interference

Double stranded RNA (dsRNA) for parental RNA interference was synthesized according to (15) and injected following the standard protocol from (27). Two non-overlapping fragments of *P. tepidariorum Sox21b-1* were isolated from the 1134 bp coding sequence of the gene: fragment 1 spanning 549 bp and fragment 2 covering 550 bp. Double stranded RNA for *P. tepidariorum Dl* (853 bp), *Wnt8* (714 bp) and the coding sequence of GFP (720 bp) as used previously (27), were transcribed using the same method. Synthesis of double stranded RNA was performed using the MegaScript T7 transcription kit (Invitrogen). After purification the dsRNA transcripts were annealed in a water bath starting at 95°C and slowly cooled down to room temperature. dsRNA was injected at 2.0 μg/μl in the opisthosoma of adult females every two days, with a total of five injections (n = 7 for each dsRNA; n= 2 for GFP controls). The injected spiders were mated after the second injection and embryos from injected spiders were fixed for gene expression and phenotypic analyses at three different time points: stage 4 (cumulus formation), stage 5 late (germ disc with migrating cumulus) and stage 9 (head and limbs bud formation).

## Supplemental Information

Supplemental Information includes five figures and two tables.

## Declaration of Interest

The authors declare no competing interests.

## Author Contributions

CLBP, SR and APM designed the project for the paper. CLBP and AS performed most of the experiments. DJL and SR carried out the genomic and bioinformatic analysis. CLBP, SR and APM wrote the manuscript with the help of DJL and AS.

## Acknowledgements

This research was funded by a CNPq scholarship to CLBP (234586/2014-1), a grant from The Leverhulme Trust (RPG-2016-234) to APM and AS, and in part by a BBSRC grant (BB/N007069/1) to SR.

---

**Supplementary Figure S1.**
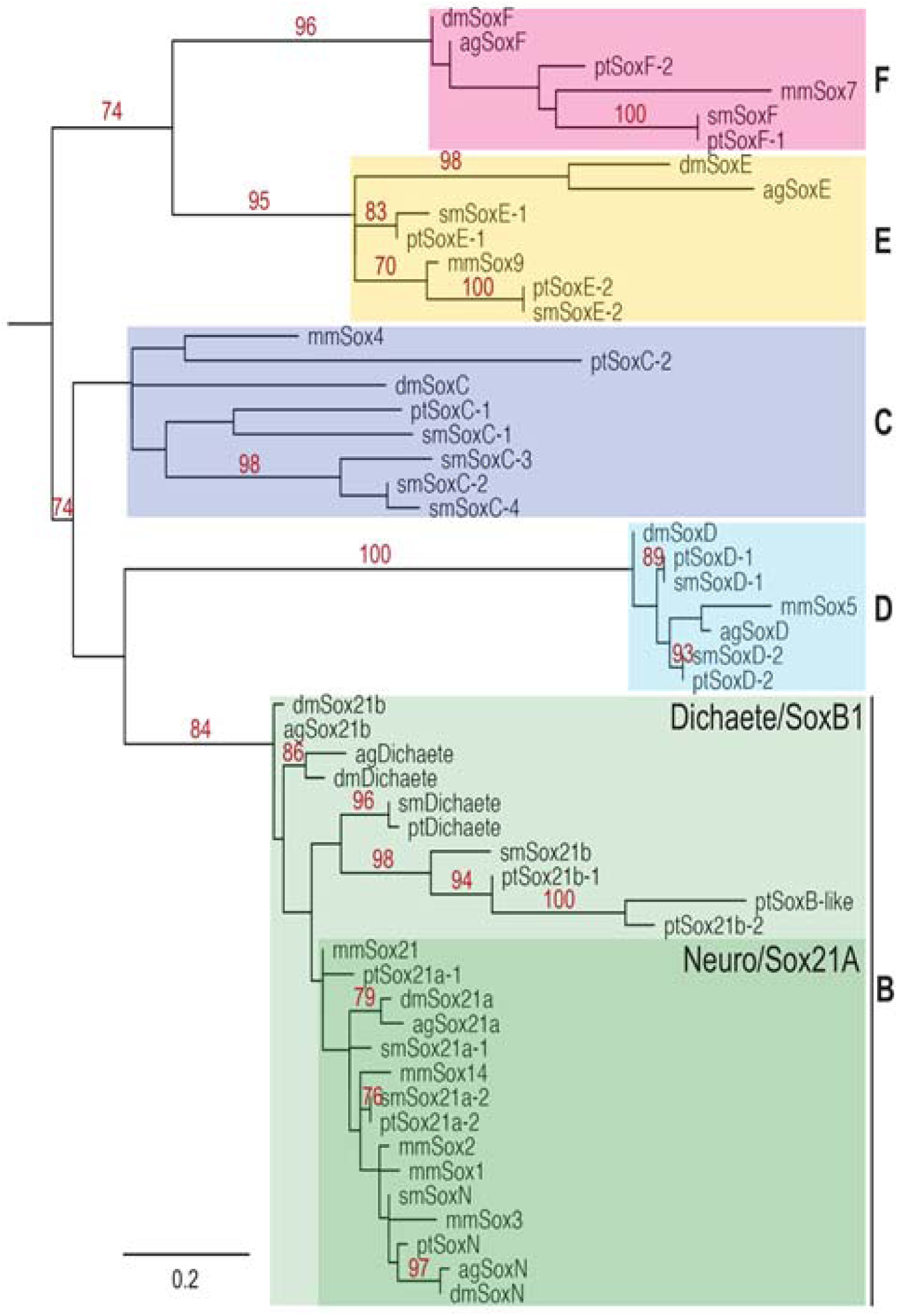
RAxML phylogeny of eumetazoan Sox genes. Phylogenetic tree made with RAxML algorithm showing the relationship between *Anopheles gambiae* (Ag), *Mus musculus* (Mm), *D. melanogaster* (Dm), *P. tepidariorum* (Pt) and *S. mimosarum* (Sm) Sox proteins based on HMG domain sequences.

**Supplementary Figure S2.**
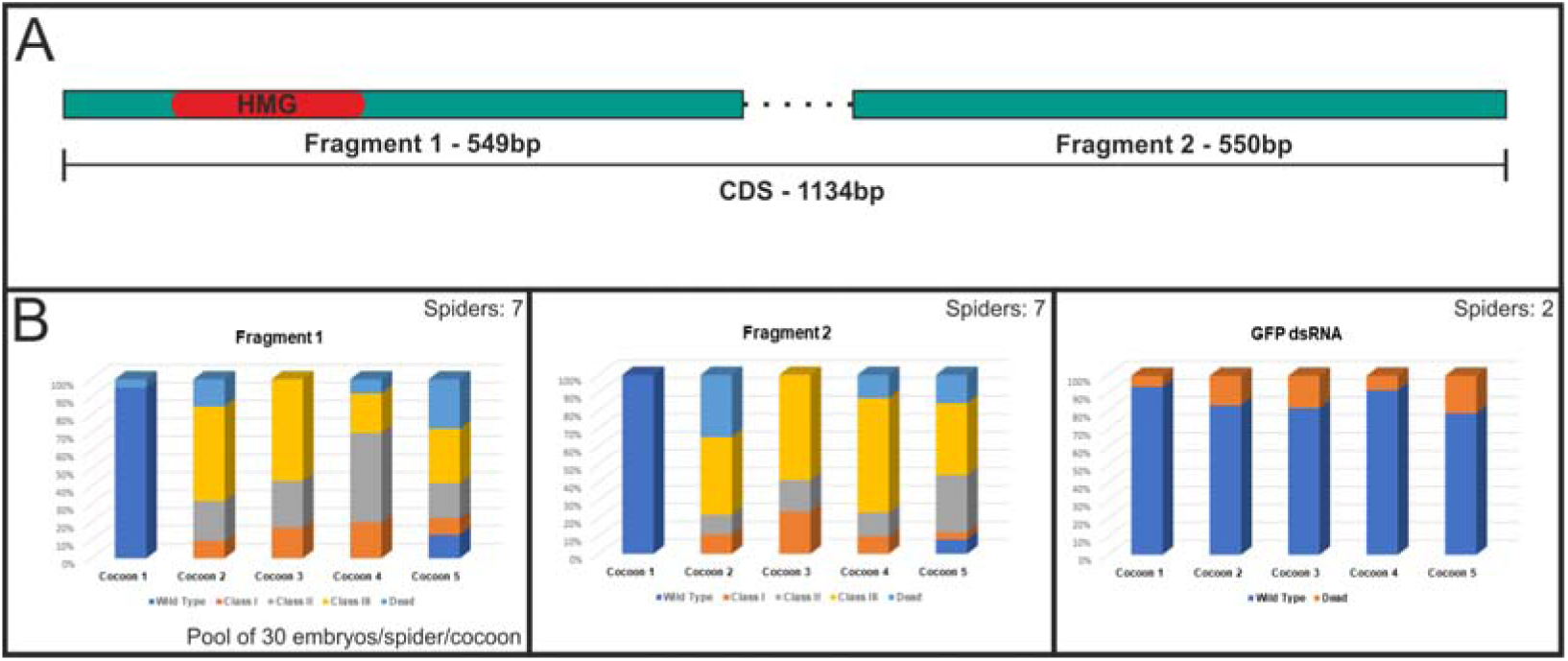
dsRNA design and phenotypical class frequencies for each fragment and GFP control injections. **A**) Two non-overlapping fragments were designed for the *Sox21b-1* coding sequence. Fragment 1 contains the HMG conserved domain (549 bp) and fragment 2 has no conserved domains (550 bp). **B**) Frequencies for each fragment, cocoon number and phenotype class. Seven spiders were injected for each *Pt-Sox21b-1* fragment and two spiders for the GFP dsRNA controls. For the phenotypical class frequencies, 30 embryos per spider per cocoon were pooled, DAPI stained and analysed (total n= 210 for each).

**Supplementary Figure S3.**
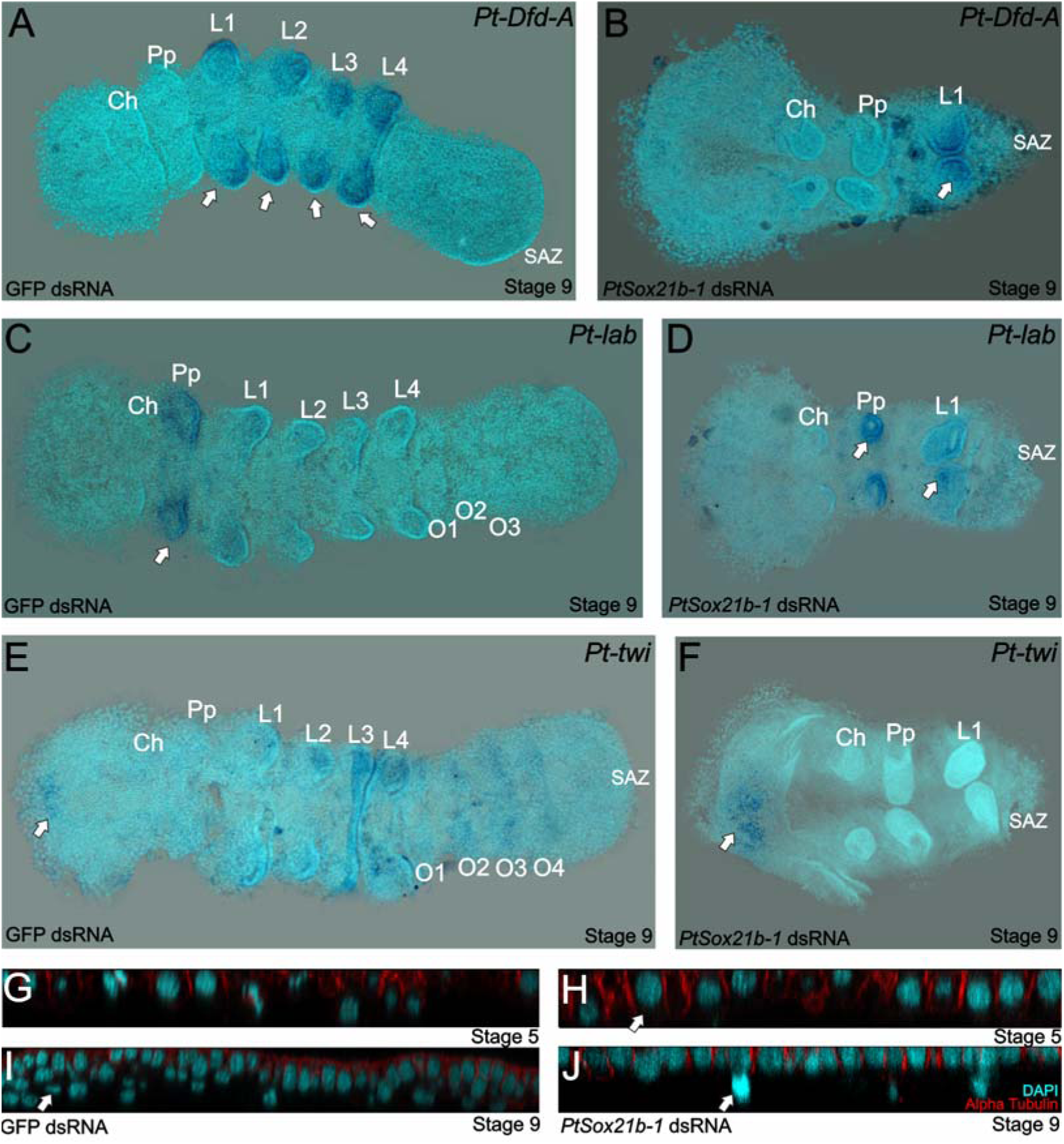
Homeotic and mesodermal gene expression at stage 9 in *Sox21b-1* pRNAi embryos. **A)** Ventral view showing *Pt-Dfd-A* expression in the limb buds of L1 to L4 segments in the control embryos (white arrows). **B**) Expression of *Pt-Dfd-A* is also observed in L1 in *Sox21b-1* pRNAi embryos (n = 9) (white arrow in **B**). **C**) *Pt-lab* is expressed in the pedipalpal segment and faintly in L1 segment in control embryos (white arrow in **D**). In *Sox21b-1* pRNAi embryos, *Pt-lab* expression can still be observed in the pedipalpal and L1 segments (n = 10) (white arrows in **D**). **E**) The mesodermal marker *Pt-twi* is expressed in the anterior-most medial region of the head, limb buds of L1 to L4, and with a stripped pattern in the O1 to O4 segments. **F**) In *Sox21b-1* knockdown embryos, only the head expression is maintained (n= 14) (white arrow in **F**). **G-J** show orthogonal projections of the cumulus (stage 5) and the head (stage 9) at 40x magnification of whole mount control embryos (left panels) and *Sox21b-1* knockdown embryos (right panels), respectively. In control embryos the formation of subectodermal layers are visible, which are lost in the knockdown embryos. DAPI stained nuclei are shown in cyan and the membrane marker alpha-Tubulin in red. Anterior is to the left in all panels.

**Supplementary Figure S4.**
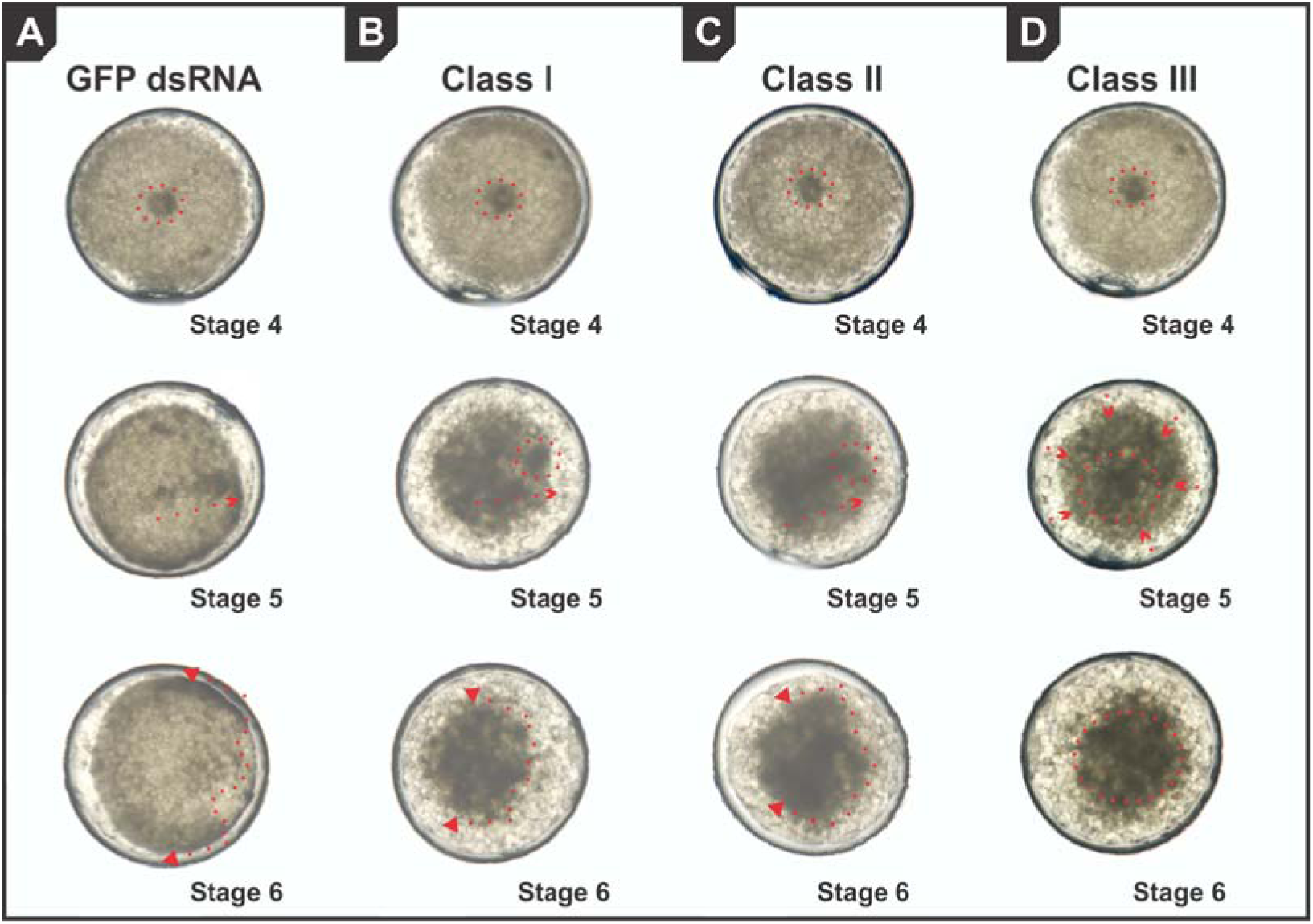
Snapshots from live imaged videos in control and *Sox21b-1* knockdown embryos. Ventral view of the germ disc in **A**) GFP dsRNA control embryos, showing the cumulus formation (red dotted lines), cumulus migration (red dotted arrow) and dorsal field opening (red dotted line and arrows). **B**) Class I *Sox21b-1* knockdown embryos showing cumulus formation, the partial migration of mesenchymal cells and limited dorsal field opening, which is also seen but more severely disrupted in Class II embryos (**C**), and absent in class III (**D**). Anterior is to the left, opposite to cumulus migration.

**Supplementary Figure S5.**
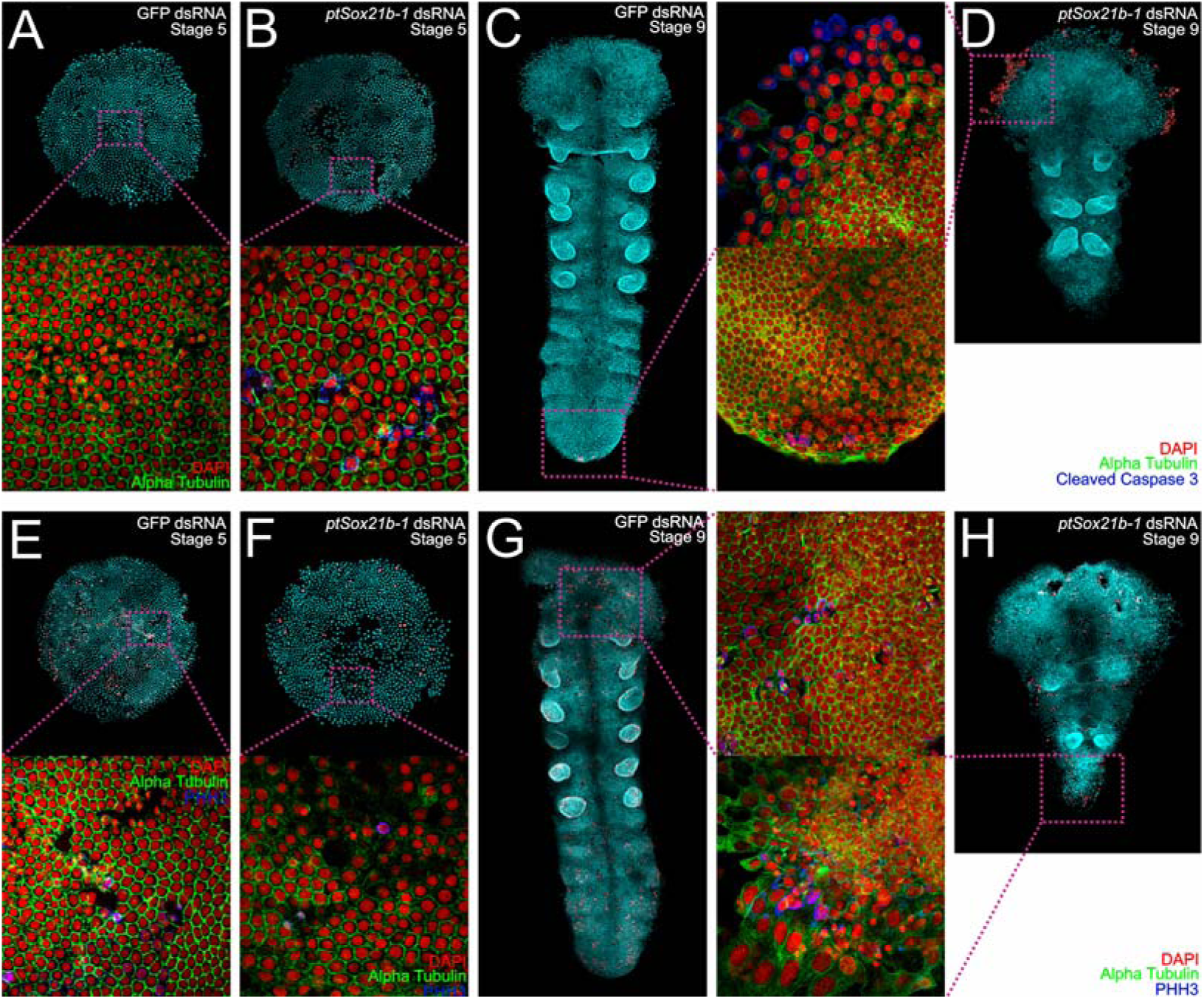
Cell death and cell proliferation in *Sox21b-1* knockdown embryos. Ventral view of stage 5 control embryos stained for Cleaved-Caspase3 (**A**) and PHH3 (**E**). Cell death is not detectable in control embryos, but a high level of proliferation can be seen. In *Sox21b-1* knockdown embryos, clusters of cells undergoing cell death can be found (**B**), as well as a decrease in proliferation in the knockdown embryos compared to controls (n= 15 for each staining) (**F**). Embryos at stage 9 stained for Cleaved-Caspase 3 (**C**) and PHH3 (**G**) show that only a small amount of cell death occurs in the SAZ, and that there is proliferation detectable throughout the entire embryo. Cell death is visible in the head extraembryonic layer in *Sox21b-1* pRNAi embryos (**D**), and less proliferation is detected in stage 9 knockdown embryos (n= 15 for each staining). Anterior is to the top. Magnifications are 100X and 400x respectively.

**Table S1.**
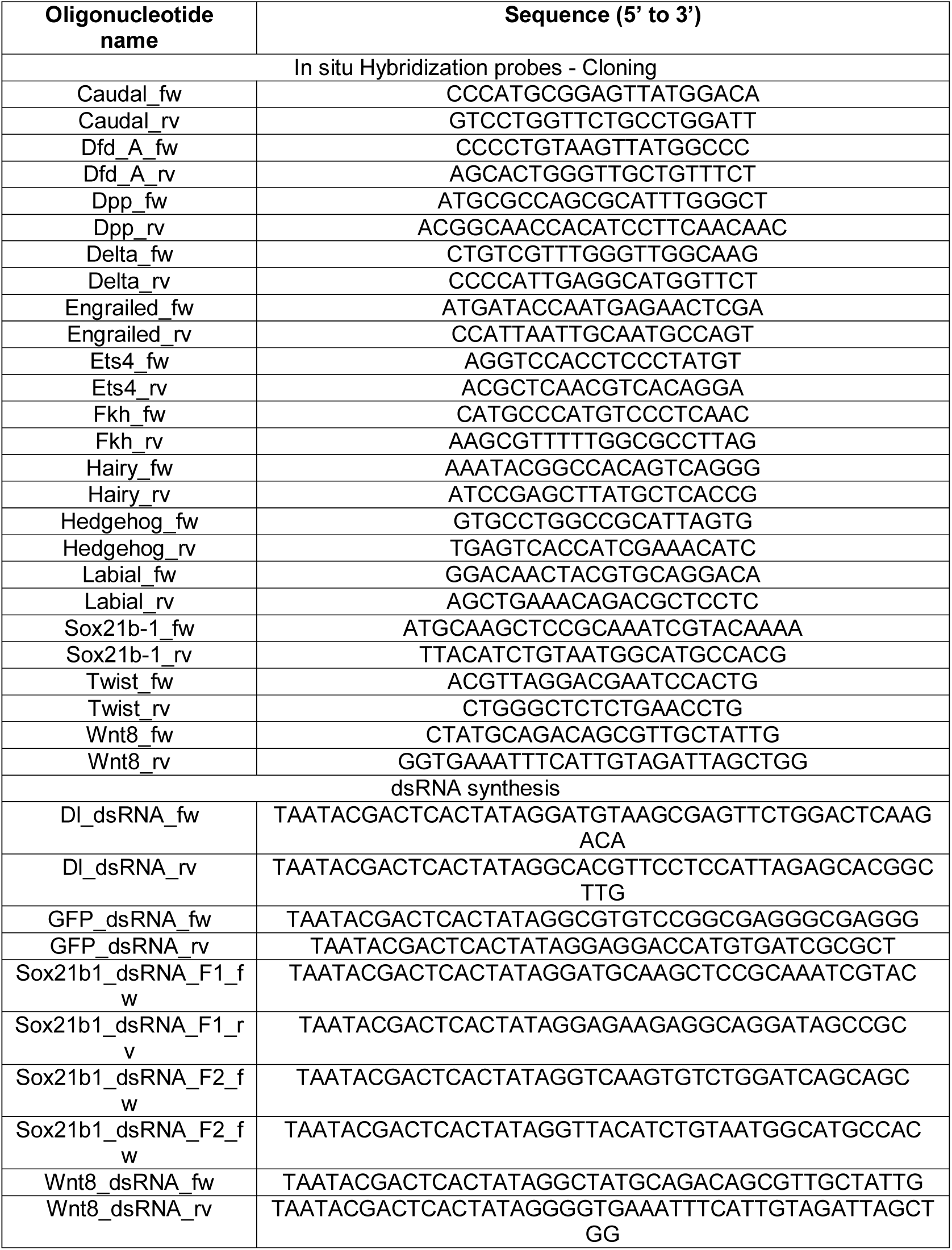
List of primers used in this paper.

**Table S2.**
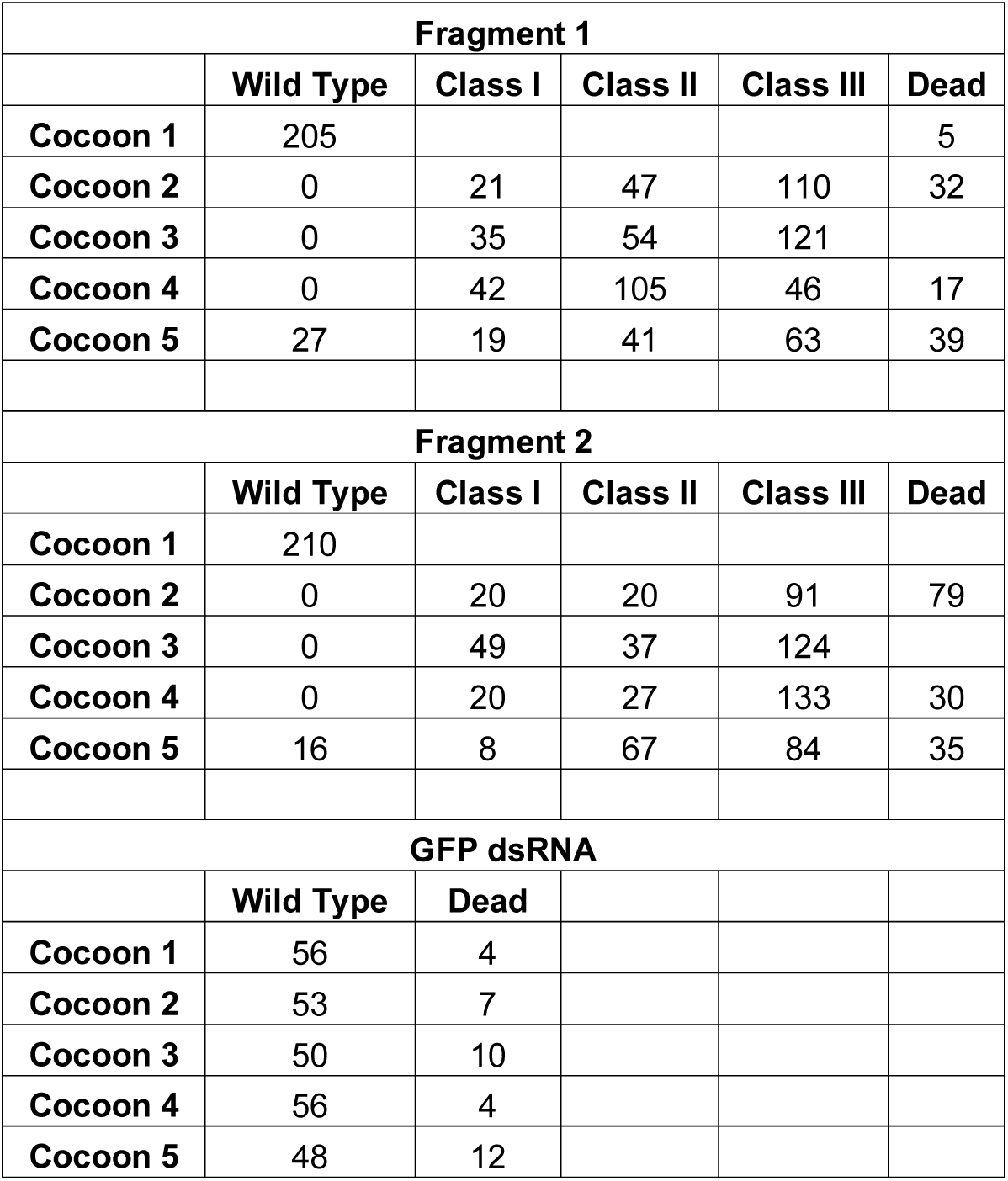
Phenotypic frequencies for each fragment (1 and 2) and GFP (control) dsRNA – 30 embryos per cocoon for each spider were pooled and the characteristics were divided into three phenotypical classes.

